# Morphogenetic processes as data: Quantitative structure in the *Drosophila* eye imaginal disc

**DOI:** 10.1101/395640

**Authors:** Bradly Alicea, Thomas E. Portegys, Diana Gordon, Richard Gordon

**Affiliations:** Orthogonal Research and Education Laboratory, Champaign, IL USA; OpenWorm Foundation, Boston, MA USA; 58 W. Grant Street #121, Healdsburg CA USA; Gulf Specimen Marine Laboratory & Aquarium, 222 Clark Drive, Panacea, FL USA; C.S. Mott Center for Human Growth & Development, Department of Obstetrics & Gynecology, Wayne State University, Detroit MI USA

## Abstract

We can improve our understanding of biological processes through the use of computational and mathematical modeling. One such morphogenetic process (ommatidia formation in the *Drosophila* eye imaginal disc) provides us with an opportunity to demonstrate the power of this approach. We use a high-resolution image that catches the spatially- and temporally-dependent process of ommatidia formation in the act. This image is converted to quantitative measures and models that provide us with new information about the dynamics and geometry of this process. We approach this by addressing three computational hypotheses, and provide a publicly-available repository containing data and images for further analysis. Potential spatial patterns in the morphogenetic furrow and ommatidia are summarized, while the ommatidia cells are projected to a spherical map in order to identify higher-level spatiotemporal features. In the conclusion, we discuss the implications of our approach and findings for developmental complexity and biological theory.

## Introduction

To advance the development and use of computational representations and models in developmental neuroscience, we require a well-characterized biological system that yields fairly unambiguous information regarding the differentiation process. We propose that differentiation of the eye of the fruit fly (*Drosophila melanogaster*) is such a candidate system (Figure 1). In the *Drosophila* eye imaginal disc (Figure 2, Supplemental File 1), ommatidia differentiation proceeds from posterior to anterior (left to right in our figures), and commences at the beginning of the third instar. It is an autoregulatory process that relies upon a complex network of molecular signals (Roignant and Treisman, 2009) triggered by a traveling induction wave (morphogenetic furrow), which has previously been identified as a wave with an alternating pattern of differentiation (Heberlein and Moses, 1995). This variety of differentiation wave (Gordon and Gordon, 2016; Gordon, 1999) is concurrent with the morphogenetic furrow and acts to control both the timing and positions of differentiated cells.

**Figure 1.**
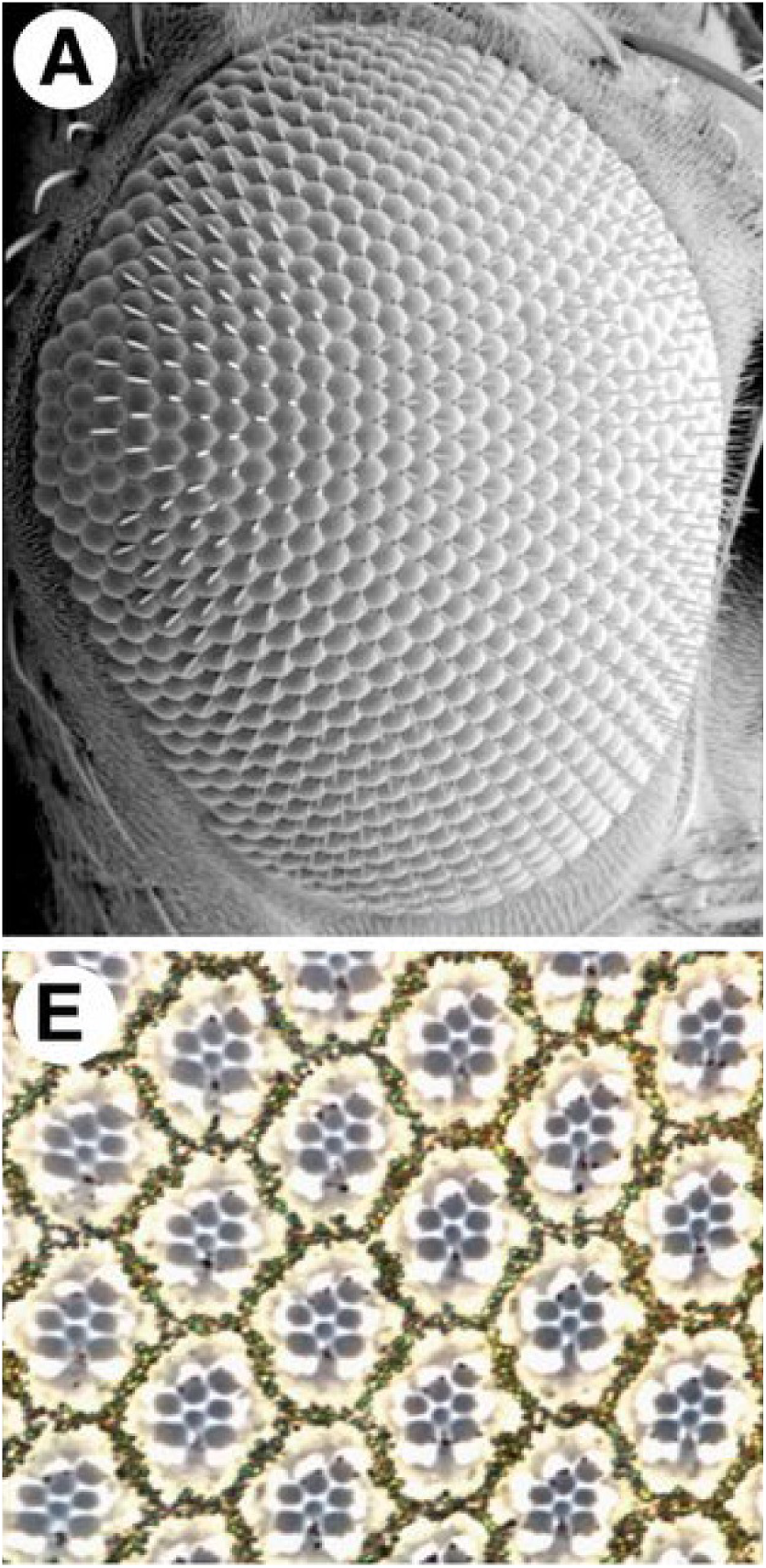
**Top:** An SEM image of the adult wild type (WT) eye in *Drosophila melanogaster*. Bottom: “The apical section of a WT eye (E) shows the typical trapezoid arrangement of rhabdomeres marking the six outer photoreceptor cells as well as the R7 photoreceptor cell (smaller centrally located rhabdomere). Granules produced by pigment cells can also be seen surrounding individual ommatidia.… Anterior is to the right.” From Baril et al. (2014) with kind permission of the Genetics Society of America.

**Figure 2.**
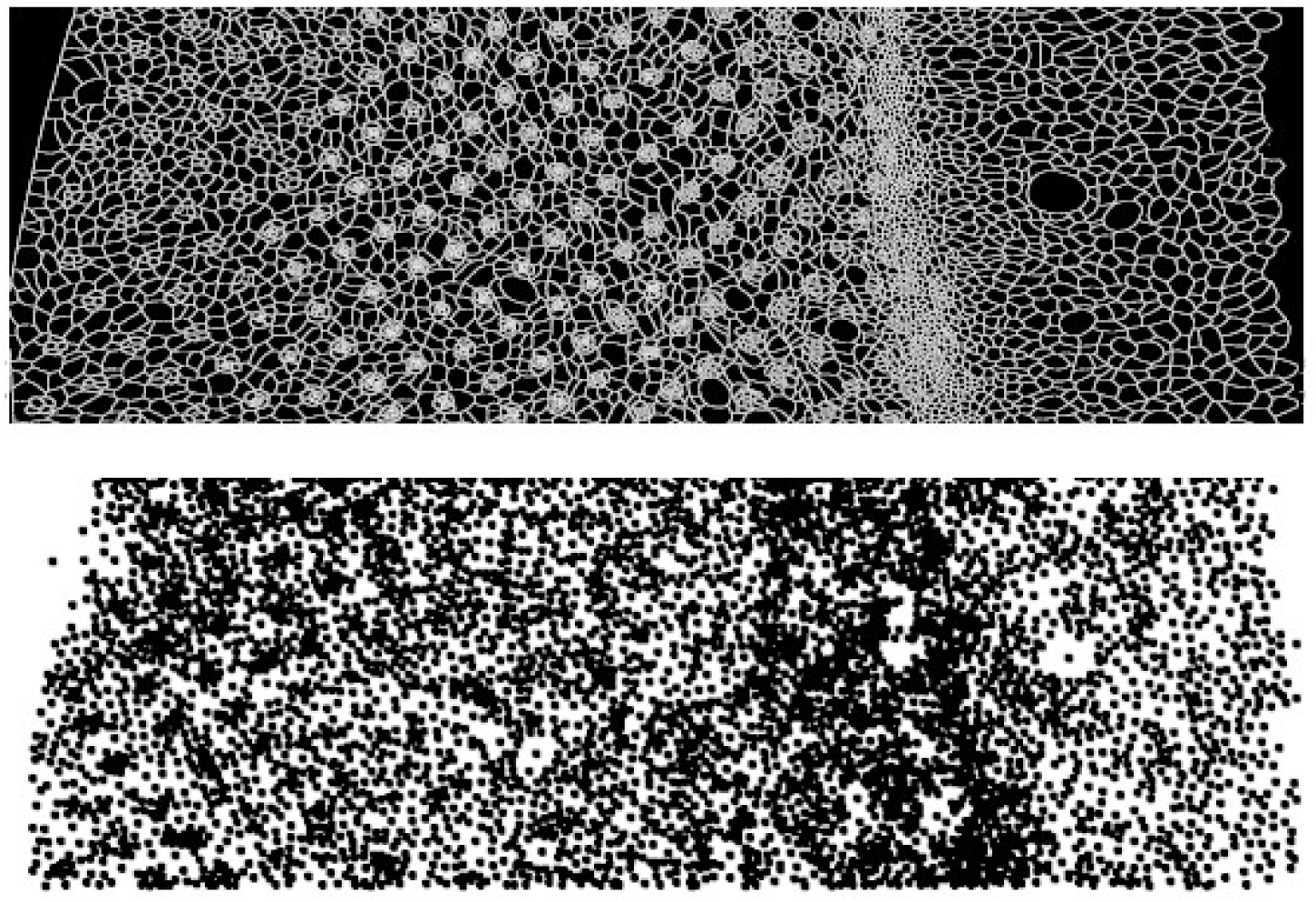
Diagram showing movement of the morphogenetic furrow from posterior (left) to anterior (right) end of the *Drosophila* eye imaginal disc. **Top:** a strip from our corrected high-resolution drawing (Supplemental File 1). **Bottom:** centroids of the cells marked by black dots. Centroids do not map perfectly to cell bodies due to the ambiguity of cell boundaries with respect to the segmentation algorithm.

In the eye imaginal disc, we seek to understand the differentiation process relative to the morphogenetic furrow. The morphogenetic furrow is a structure that is defined by apical constriction and apical-basal contraction (Gordon and Gordon, 2016; Lee and Treisman, 2002; Schlichting and Dahmann, 2008). This furrow produces 800 ommatidia structures present in the adult compound eye by inducing proneural states during its movement through undifferentiated cells (Chanut and Heberlein, 1995). The cells in this epithelium commit to a neuronal fate as they receive signals triggered by the passing of the morphogenetic furrow and its proximity to/recruitment by an ommatidia founder (R8) cell (Brennan and Moses, 2000; Dokucu et al., 1996). Yet not all cells in this sheet commit to a neuronal fate (Wolff and Ready, 1991), and we characterize these as “background cells”. While quite regular, this process also reveals a degree of intrinsic variation (Swain et al., 2002) between eye imaginal discs. Various molecular pathways interact with progression of the furrow during differentiation of various cells in a single ommatidium (Davis and Rebay, 2018; Greenwood and Struhl, 1999). These patterns of differentiation may be due to a phenomenon we have defined as single-cell differentiation waves (Gordon and Gordon, 2016; Gordon, 1999).

Our data consists of a single, high-resolution camera obscura drawing (Supplemental File 1) of a *Drosophila melanogaster* eye imaginal disc observed during the third instar of development (Wolff, 1993). While eye imaginal discs are highly-stereotyped in terms of axial patterning (e.g. a lattice of ommatidia), there also exists variation both within and between organisms (e.g. positional and biological noise) in terms of the positioning and fate of specific cells within the disc (Heberlein and Moses, 1995; Tare et al., 2013) (for a definition of biological noise, see Elowitz et al. (2002)). Therefore, while these data are representative, they by no means capture the variation inherent in the differentiation process. These data are unique in that the imaginal disc has been caught in the act of differentiating with every cell recorded (Figure 2, Supplemental File 1). A morphogenetic furrow marks the boundary between a population of isotropic and presumably undifferentiated cells to a structured population of ommatidia cells and background cells. Here we use both mathematical and computational techniques to uncover patterns, features, and geometric relationships previously not characterized in the literature.

The analyses here also provides a basis for future studies using cell segmentation, cell tracking, and lineage tracing (Meijering et al., 2009) to produce a dynamic view of the process. This type of systems morphometrics has the potential to inform molecular investigations as well as computer simulations of the developing *Drosophila* eye (Formosa-Jordan et al., 2012). We will also make the case that computational analysis of ommatidia morphogenesis is deserving of more sophisticated time lapse microscopy, well beyond the limitations of a camera lucida-based drawing.

The current study is uniquely positioned in the literature. By focusing on the analysis of cell position in a static image, we provide a top-down perspective missing in the contemporary literature. Most previous studies applying morphometric approaches to the *Drosophila* imaginal disc have focused on two topics: growth of the imaginal disc proper and presumed morphogen gradients. Vollmer et al. (2016) and Bittig et al. (2008) have introduced quantitative models of imaginal disc growth in terms of cell number and tissue mechanics, respectively. In the former study (Vollmer et al., 2016), it was found that growth of the imaginal disc structure is independent of cell number, so that variation in cell number has no effect on the size of the disc. This is analogous to the independence of urodele amphibian size with cell number, which can be manipulated by varying polyploidy (Gordon and Gordon, 2016). The latter *Drosophila* study (Bittig et al., 2008) accounts for the anisotropy of cell division across the natural variation of imaginal disc formation.

There are also a number of studies focusing on the relationship between cellular differentiation at the morphogenetic furrow and gene expression gradients. Fried et al. (2016) and Fried and Iber (2014) introduce a parametric model of regulatory interactions relative to movement of the morphogenetic furrow. The gradients that result from various molecular interactions tend to predict the speed of morphogenetic migration across the disc (Fried et al., 2016). Similarly, (Garcia et al., 2013) use a linear regression model to predict gene expression gradient boundaries in the context of imaginal disc anatomy.

A number of studies have modeled morphogenesis as a kinetic phenomenon. Kicheva et al. (2012) and Lander et al. (2002) have approached morphogenesis as a problem of molecular diffusion. Lander et al. (2002) introduce a set of differential equations for modeling the diffusion of various molecular factors across the imaginal disc with respect to morphogenetic furrow position. Kicheva et al. (2012) expand on this model by using a parametric approach that includes parameters for production and degradation as well as diffusion. Averbukh et al. (2014) expand on this line of work further by introducing an expansion-repression feedback loop, which specifies how and when cells are exposed to the expression and repression of various molecular signals.

In addition to applying a series of quantitative techniques, we also wish to establish an open dataset as well as to extract information about a single developmental process. A previous analysis of these data only estimated one visually invisible, global pattern, an alternation between two large and two small cells along the bottom of the furrow using a variogram method (Gordon, 1999). Our approach here is much more comprehensive, and applies image processing techniques, feature selection methods, and geometrical transformations to the same source material. This paper presents an exploratory analysis of the data along with four computational hypotheses, the latter of which may ultimately lead to both local and global statistical invariants.

### Computational Hypotheses

We propose four computational hypotheses that can be realized through analysis and transformation of the quantitative data. We will address these hypotheses using a variety of methods. These can be stated as follows:

**H1:** the size distributions of cells representing three components of the eye imaginal disc (furrow, differentiated background, and ommatidia) will yield differences.
**H2:** characterizing ommatidia complexity (size and position) can be informative for understanding the tempo of differentiation in both space and time.
**H3:** projecting ommatidia to a spherical map can provide a uniform representation on which to model the process of differentiation.
**H4:** animations of the spherical map as a series of time slices will reveal a process analogous to anatomical differentiation.

## Methods

The poster (Wolff, 1993) accompanying Wolff and Ready (1993) was digitized and the loops, representing cell boundaries, were closed digitally by hand, where needed (Figure 3), as shown in Supplemental File 1. By comparing our digitized image showing about 150 pixels between ommatidia with Figure 11 in Wolff and Ready (1993), with an average distance between adjacent ommatidia of 7.2 μm, we estimate the width of our pixels at 0.05 μm. The image was then inverted and segmented using the ImageJ (Rasband, 2018) particle analysis function. This resulted in discrete cells (n=9733) each with a defined area and two-dimensional centroid position. The areas are projections parallel to the microscope axis. We have not attempted to correct for curvature, which requires accurate cross sections (Baker, 2007; Bessa et al., 2002; Gibson et al., 2002).

**Figure 3.**
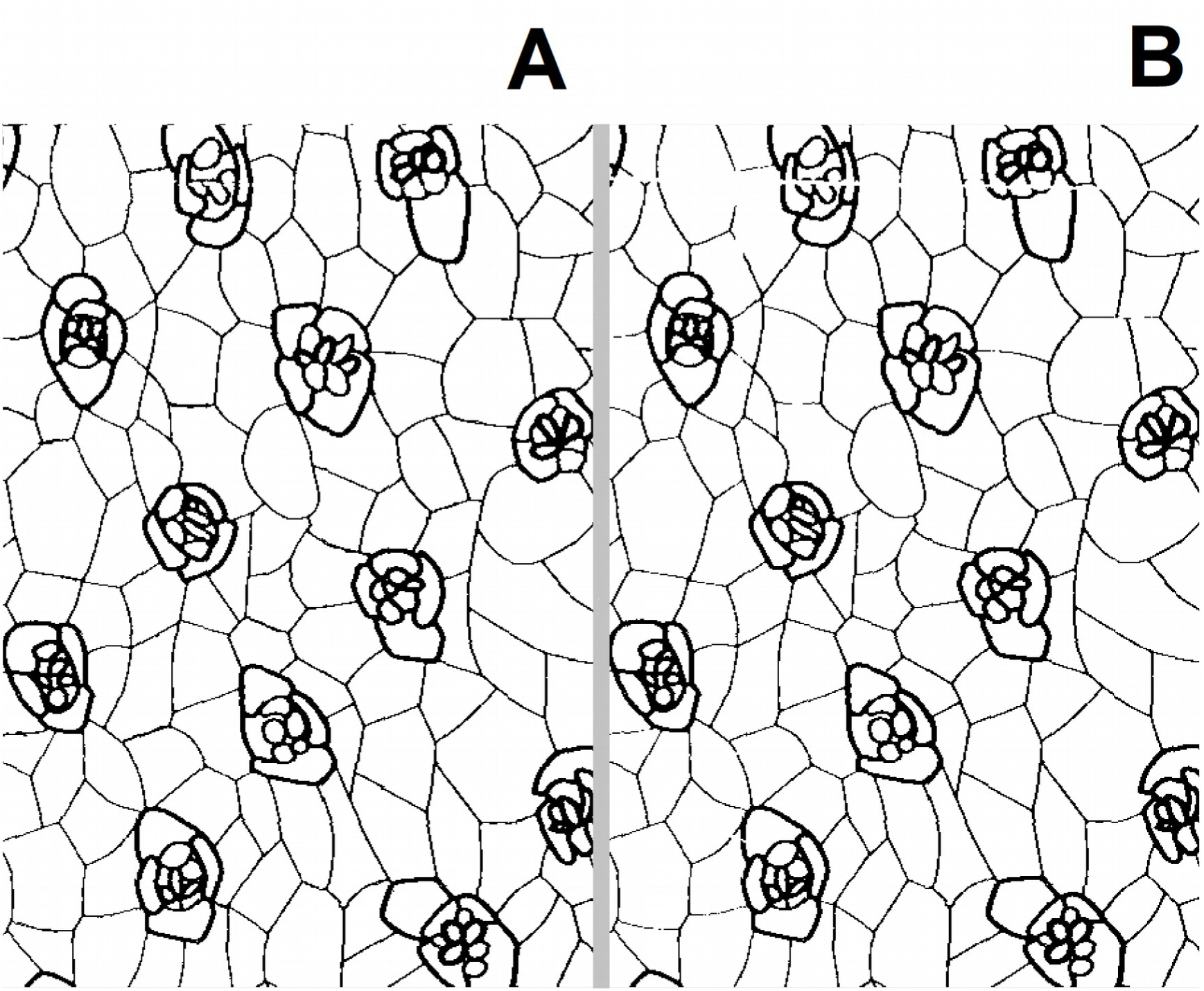
Closeup of the camera obscura sketch of all cells in a *Drosophila* imaginal disc (Wolff, 1993). **A:** before correction. **B:** after closing by hand all cell-cell boundaries that had gaps. For the full, corrected sketch see Supplemental File 1. Note that the original sketch had thick lines designating ommatidial cells.

Inspection of the inverted image suggested that all cells were segmented from one another. The segmented image was further transformed using two types of transformation: spherical projection and realignment. Realignment was achieved by using a linear regression function to align the image about the central region of the morphogenetic furrow identified by cell density and the ends of ommatidia rows defined through a ridge estimation. The retouched image was also decomposed manually into layers containing: 1) cells associated with the ommatidia, 2) cells serving as the background to the ommatidia and anterior to the furrow, and 3) cells associated with the furrow. Layers were defined by transitions in pattern, cell size and line thickness.

### Open Data and Analysis

We have provided extensive documentation of our data and analysis in a version-controlled repository. Anyone can become a collaborator, download data and methods, and contribute to the analysis. All code, analyses, and associated images are publicly available on Github (Alicea, 2018a).

### Binary Maps

Binary maps were created and are in the publicly-available repository. All images were reduced to 1-bit binary images and decomposed into a numeric matrix, with pixels labeled “1” being cells of interest and pixels labeled “0” being background cells. Ommatidia and background layers were decomposed into binary maps and reconstructed using SciLab 6.0 (Paris, France) and the IPCV 1.2 toolbox (Luh, 2017). Binary maps were used to verify segmented images and their relationship in the ommatidia.

All binary maps are presented in the Github repository. Binary maps for “BG-left” and “ommatidia” were created from the “background” and “ommatidia” masks as described in the Methods section. Each mask was reduced to a 1-bit image in ImageJ, then converted into a binary matrix using SciLab 6.0. Binary images describe every pixel that belong to a cell, and were coded with a value of “1”. This binary matrix can be reduced to a set of *x,y* coordinates for which the color value equals “1” using the SciLab code in files “code-for-binary-images.md” and “ht-make-xy-points-from-binary-matrix.md”.

### Discrete Feature Spaces

To identify specific features in the segmented data, we created several layers which were then used to segment and label specific types of features. These layers included the furrow layer (n=1433), the background layer (n=3966), and the ommatidia layer (n=3811). The remaining 523 cells are prefurrow cells (Figure 4). The ommatidia layer was later limited to all cells larger than 100 pixels (n=3249). The most informative of these is the ommatidia layer, which features only cells associated with ommatidia that form multiple rows across the differentiated surface of the imaginal disc. A labeled dataset was created from the ommatidia layer and included all segmented cells over 100 pixels in size. Each cell was assigned to a cluster, which was labeled by row and order from anterior to posterior.

**Figure 4.**
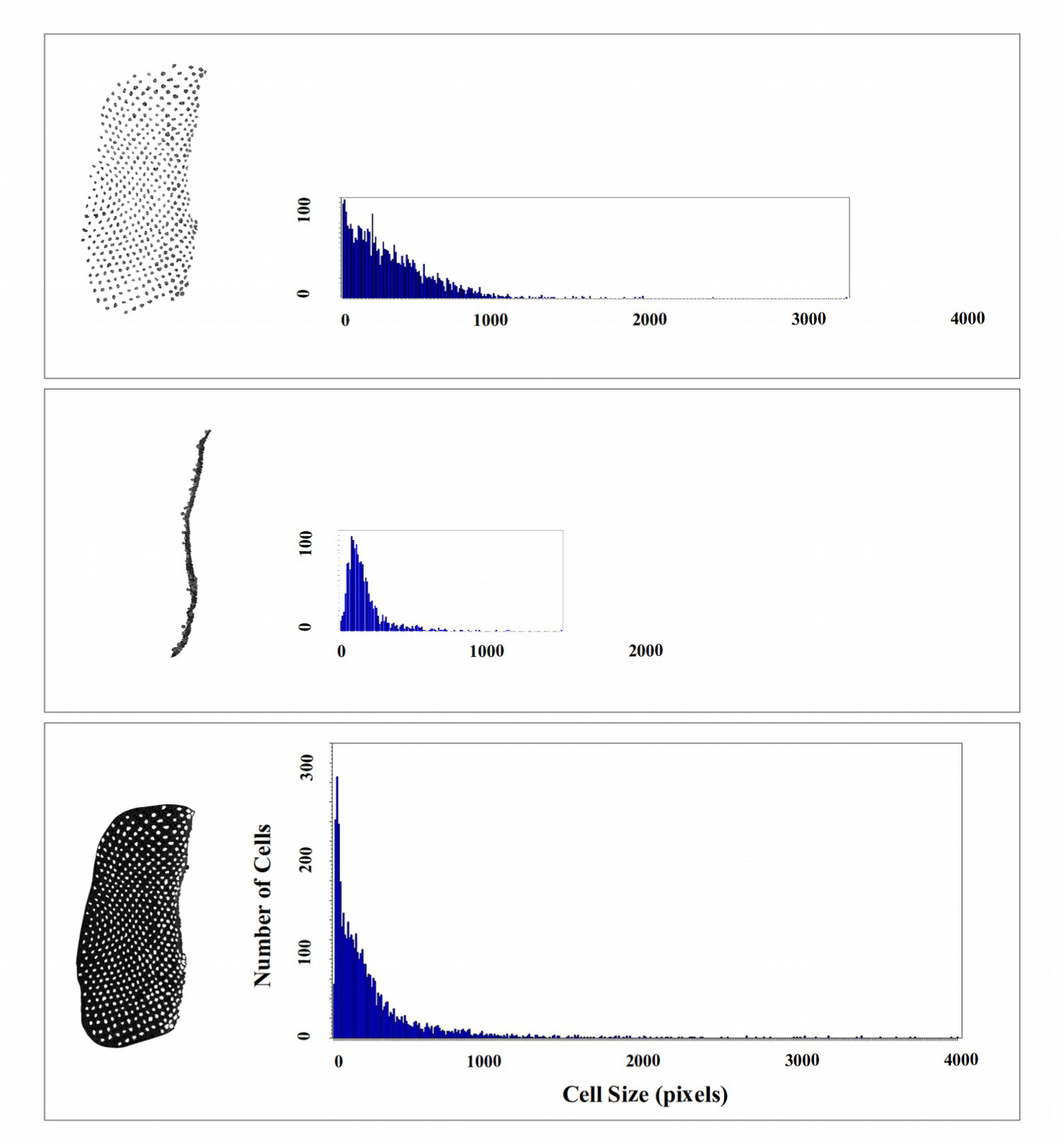
A (left): digitally isolated ommatidia. A (right): histogram summarizing the distribution of cell sizes in A (left) – (n=3249, 325 bins). B (left): isolated furrow. B (right): histogram summarizing the distribution of cell sizes in B (left) – (n=1433, 143 bins). C (left): isolated background cells. C (right): histogram summarizing the distribution of cell sizes in C (left) – (n=3966, 397 bins). The distributions have been recalled so that they can be directly compared. Cell sizes (areas) are given in terms of number of pixels, with each pixel a square 0.05 μm on a side.

### Furrow Alignment

A linear regression function was used to align and rotate the segmented imaginal disc about the furrow to represent the furrow as a straight vertical line. Using 18 candidate points estimated from center of the furrow layer, a linear regression equation was calculated. The *a* and *b* parameters were then optimized until a straight line was obtained. The optimal function is: *y* = 11.57 + 9722.8*x*. All *x* and *y* coordinates were then transformed to *x′* and *y′* using the following equations

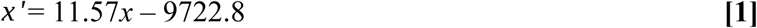

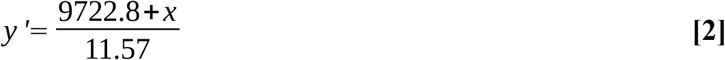

We have also developed software to straighten the furrow using a direct pixel manipulation of the raw image. The mean of all *x* coordinates of a selected set of image pixels is used to create a linear vertical target line. Pixel rows (sets of *y* coordinates) are then rectified by horizontally shifting their selected pixel to the mean. While this alternate method straightens the eye imaginal disc furrow in the raw (unsegmented image), and may be useful for exploratory or analysis of microscopy images, it also introduces waviness in the cell bodies that was not desirable for our particular set of analyses. This program is located in the public Github repository for use by the *Drosophila* community (Alicea, 2018b).

### Ridge Estimation

To determine which cell clusters constituted rows of ommatidia, ridges were estimated from a scatterplot of all cells anterior to the morphogenetic furrow. A ridge can be defined as an n^th^ order polynomial function that passes through an aligned series of cell clusters. Clusters are defined as regions of high density in the scatterplot, or regions where more than 10 centroids are fused together in the image.

### Spherical Map

The two-dimensional realigned ommatidia were mapped to a spherical representation using SciLab 6.0 (Paris, France). This provides a three-dimensional volume within which to explore the data. The dimensions of this space are defined mathematically as

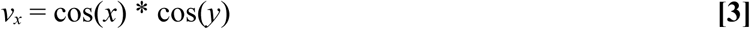

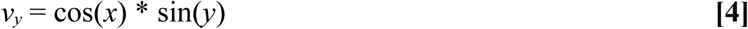

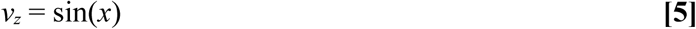

While this approach decorrelates the points in a single ommatidia with respect to spatial position, it also normalizes the sources of curvature. The range of x-axis positions in the data also represent discrete periods of time since differentiation. As one moves anteriorly across the eye imaginal disc (represented by cells with increasingly larger values in the aligned segmented images), the greater is the time since differentiation (see Figure 2). Spatial positions were taken from aligned *x,y* coordinates and then projected to the spherical map with cell size information added.

### Heat Maps

To complement the spherical maps, a heat map was created by binning the cell data from each row of ommatidia (*y)* across the non-aligned segmented image from the anterior end to the morphogenetic furrow (*x*). Each bin of 100 pixels in width (or 5 μm) represents an interval of space-time (a set of spatial positions on the eye imaginal disc surface and distance anterior from the morphogenetic furrow). The color gradient represents the number of cells (denoted by centroids) found in a specific bin.

## Analysis

### Cell Size Distributions

To get a feel for the segmented data, we constructed a rank-order frequency plot that shows a relationship many small cells and relatively fewer large cells. Figure 5 shows the data for all segmented cells in the eye imaginal disc. The graph shows not only the expected preponderance of smaller cells, but also the range of variation at smaller sizes. One notable feature of this distribution is a diversity of sizes that is consistent both among smaller and larger cells. This may be related to the independence of differentiation waves from cell size, as is apparent in polyploid salamanders (Gordon and Gordon, 2016). The cause of the wide range of cell sizes in the *Drosophila* eye imaginal disc is not known.

**Figure 5.**
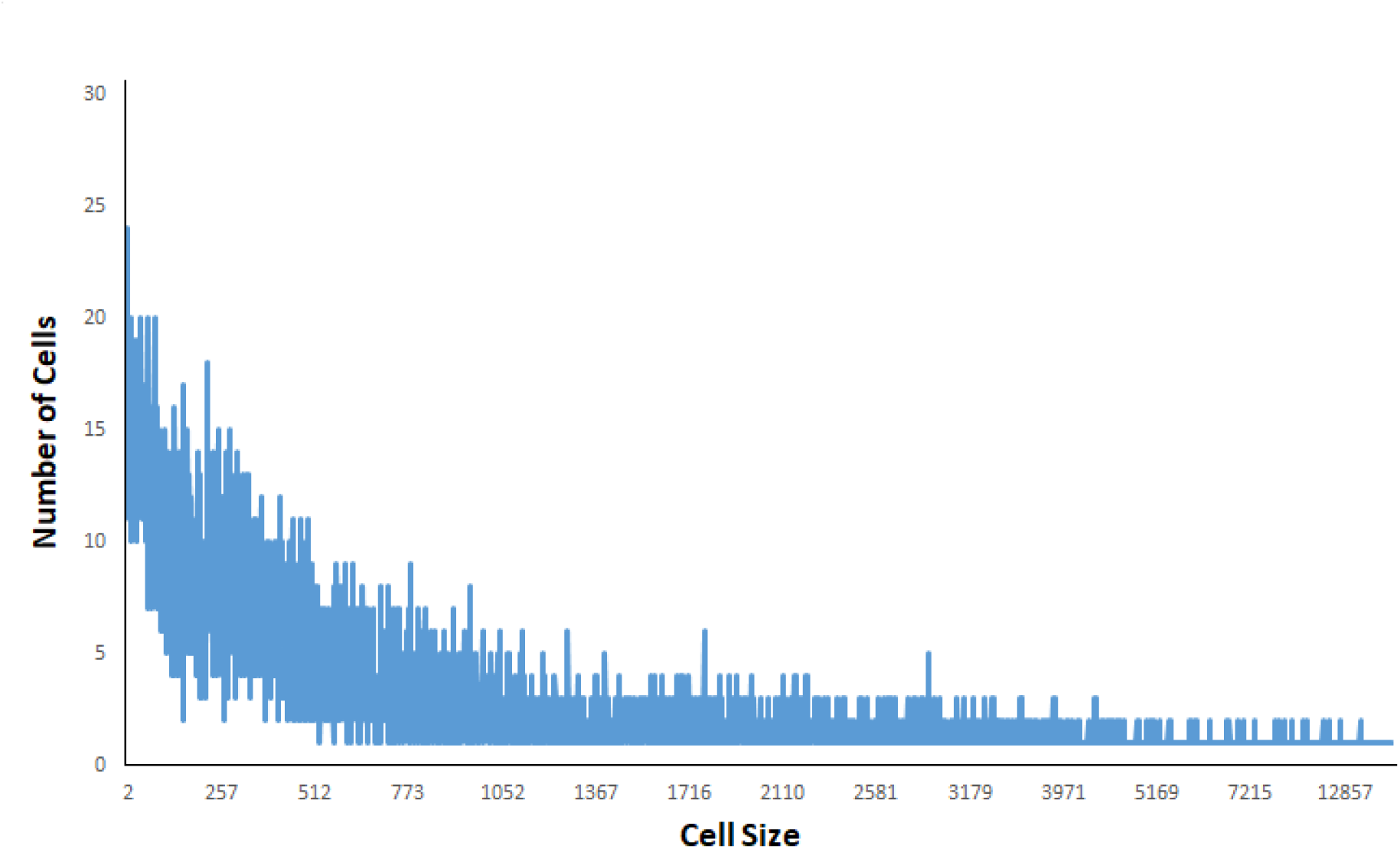
Rank order frequency plot (exponential distribution) for all cells in the non-normalized eye imaginal disc segmentation. Cell size is shown as number of pixels (projected apical area). One pixel equals 0.05 μm. All cells of size 1 and greater than size 40000 pixels were eliminated to minimize statistical artifacts. The small “cells” may be noise artifacts of the segmentation algorithm.

To follow up on the diversity signatures, we also looked at a more restricted set of regions within the eye imaginal disc. The cell size distribution of the isolated furrow, isolated background, and isolated ommatidia were calculated and compared. This was done using three histograms, and the analysis shows differences in cell size distribution between each region in shown in Figure 5. These differences are due to processes related to differentiation and each structure’s relationship to this process.

A comparison between the histograms does reveal a few trends worth noting. Cells labeled as “background” tend to have many very small constituents, while those labeled “furrow” and “ommatidia” have a bit more size diversity. Notably, the furrow has a bit more diversity at larger size scales, but in a different way than the bulk size distribution. These size distributions challenge the view of the eye imaginal disc as a “crystalline” array (Ready et al., 1976), and suggest a more complex quantitative arrangement such as found in the phyllotaxis (spiral arrangement) of sunflower seedheads (Swinton and Ochu, 2016).

### Furrow Straightening

One strategy used to correct for curvature of the furrow along its length is to find a canonical furrow and align all cells to the linear function. This is done using a linear regression function to realign the cells. A plot of the non-layered realigned eye imaginal disc and alignment function are shown in Figure 6. We can clearly identify centroids marking individual ommatidia in this plot. These centroids (representing cells of different sizes) appear as clusters against a background of individual centroids or smaller clusters. The relatively large size and alignment of these ommatidia-related clusters allows us to apply an estimation procedure to identify larger-scale features such as sequential rows of ommatidia. We use an approach called ridge estimation to make these identifications, which is similar to the serial application of polynomial regression functions.

**Figure 6.**
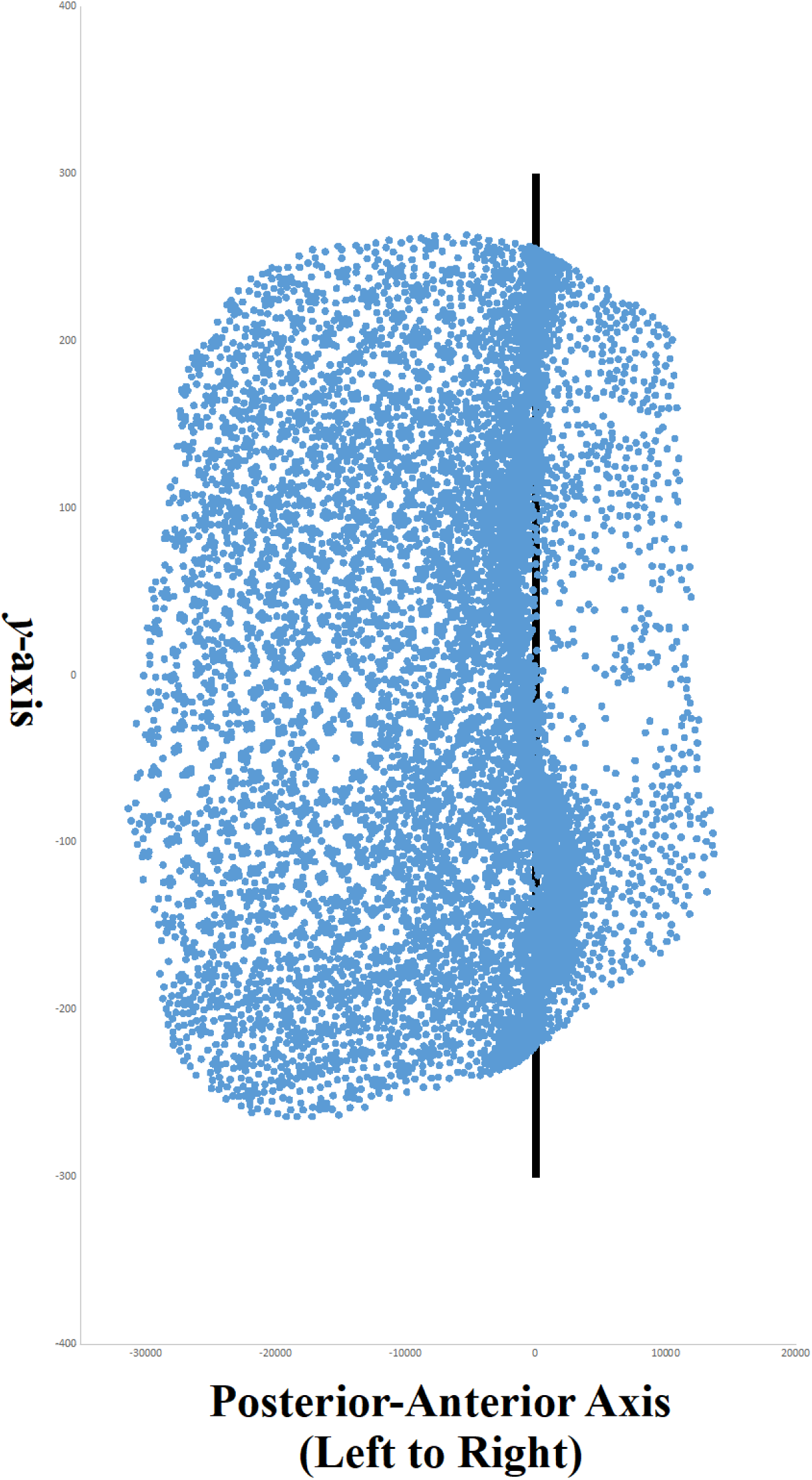
Image of the segmented *Drosophila* eye imaginal disc centered upon an vertical line that runs through the mean of the furrow region. **Blue dots:** cell centroids, **Black line:** average x-position of the bottom of the furrow.

Using all segmented cells (centroids) in the anterior (differentiated region) of the furrow region, ridges were estimated from the central tendency of all centroid clusters in the image (Figure 7). The horizontal relationships (physical continuity) between these centroid clusters were used to determine the positions of individual ommatidia (see Methods), while the ridges themselves were used to define distinct rows of ommatidia ordered from top to bottom (in descending order) along the *y* axis (for numeric values, see Supplemental File 2). This method reveals 26 rows of ommatidia, which was refined to 28 rows upon further visual inspection of the data due to positional clutter and geometric ambiguity towards the bottom of the image. These rows also define the *y* axis of the heat map shown in Figure 8.

**Figure 7.**
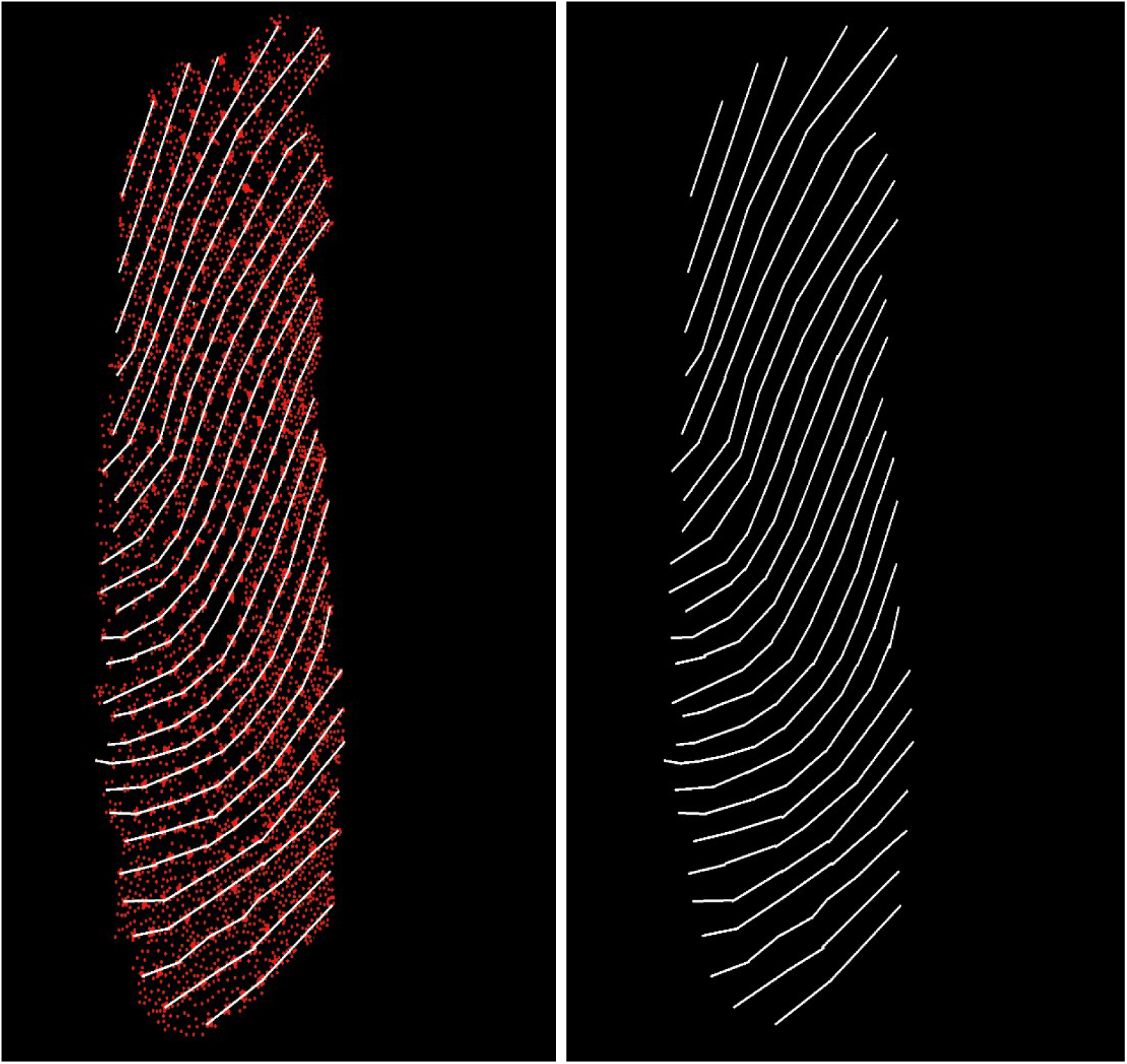
Ridge estimates that reveal 26 distinct rows of ommatidia across the differentiated section of the *Drosophila* eye imaginal disc. **LEFT:** ridge estimation from segmented centroids. **RIGHT:** ridges in isolation demonstrating the estimated contours of each row.

**Figure 8.**
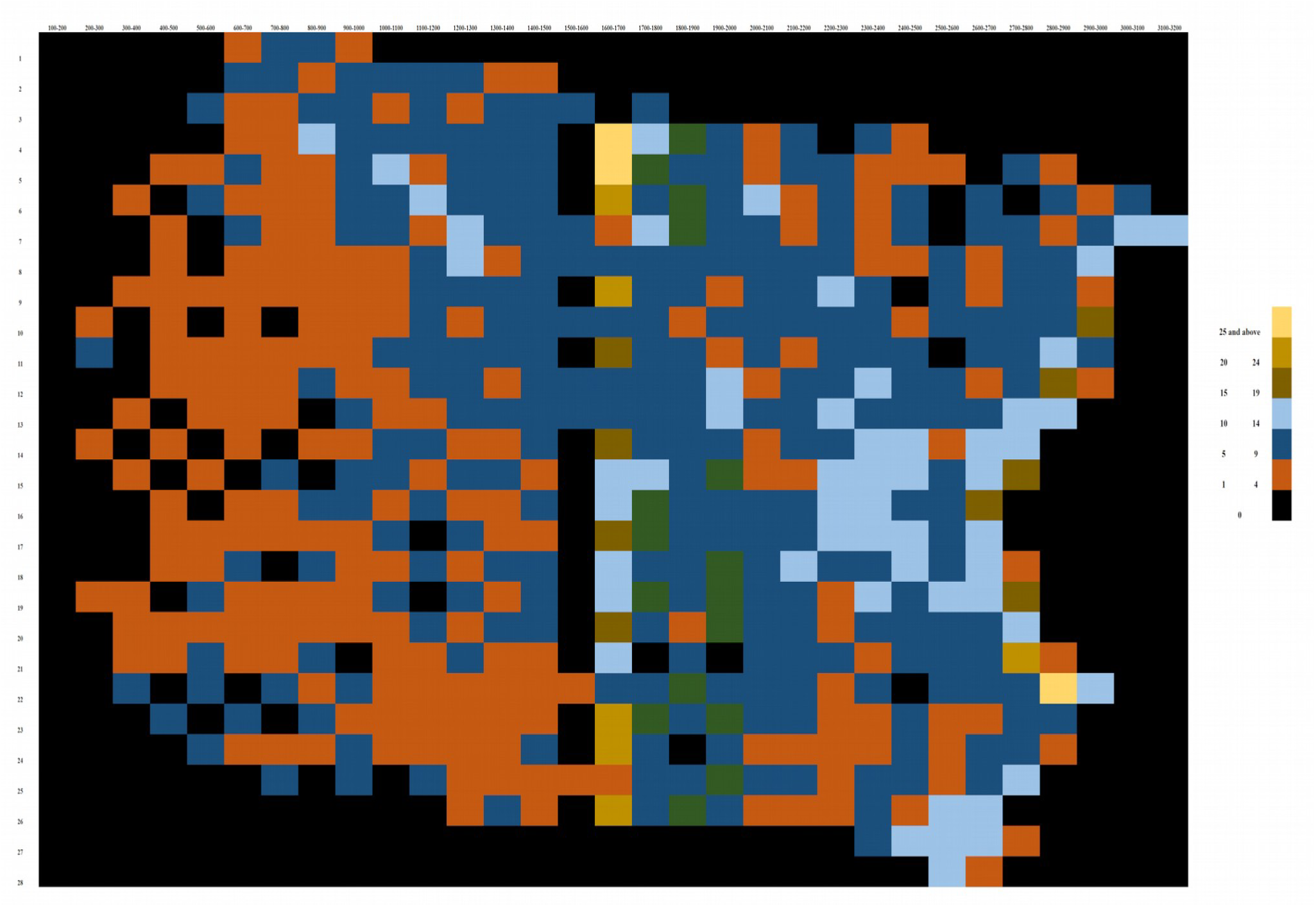
Heat map that decomposes the eye imaginal disc into 31 space-time slices (sampled at 100 pixels per bin) across 28 rows of ommatidia (sorted from top to bottom of image. The heat map was reconstructed from the non-aligned disc reconstructed as a series of straight rows. One pixel equals 0.05μm. Anatomically, the heat map extends from the anterior end of the disc (left) towards the morphogenetic furrow (right hand black edge of the colored pixels). Cells are color-coded by the number of cells in each bin.

### Ridge Estimation in the Differentiated Region

### Characterizing Ommatidia Complexity and Heat Map Decomposition

To understand the labeled ommatidia cells in more detail, we quantified individual ommatidium and compared them across individual rows and columns. Rows and columns were defined by their cluster and ridge position, respectively. A statistical summary of each ommatidium (a distinct *row*, *order* position) and its component cells is presented in Supplemental File 2. These data can also be used to stratify the spherical map project and show the locations of specific ommatidia and their component cells.

Based on this statistical summary, a heat map representing the differentiated portion of the eye imaginal disc as a two-dimensional matrix is shown in Figure 8. Each row is shown as a straight column, while 100 pixel (5μm) slices of the anterior-posterior axis serve as individual rows. The heat map reveals a two-dimensional frequency analysis of differentiated cell density. This analysis reveals an increasing density of differentiated cells (more populous ommatidia) moving from the posterior pole towards the morphogenetic furrow.

There also seems to be a streak of no cells in the middle of our sample that might be an artifact of treating each row as a straight line. On the other hand, it might mark a previously uncharacterized global event in the differentiation process. Regardless of cause, there are empty bins scattered throughout the matrix (which decrease in number as we move in an anterior direction). We suspect this may be due to a combination of inaccuracies in the drawn representation and variation in the differentiation process.

### Projections to Spherical Map

To understand higher-order patterns more similar to phyllotaxis than crystalline arrays in the ommatidia feature space, as well as to reduce the multidimensional effects of ommatidia ridge curvature (see Methods), these data were projected to a spherical coordinate system. This spherical map is shown in Figure 9.

**Figure 9.**
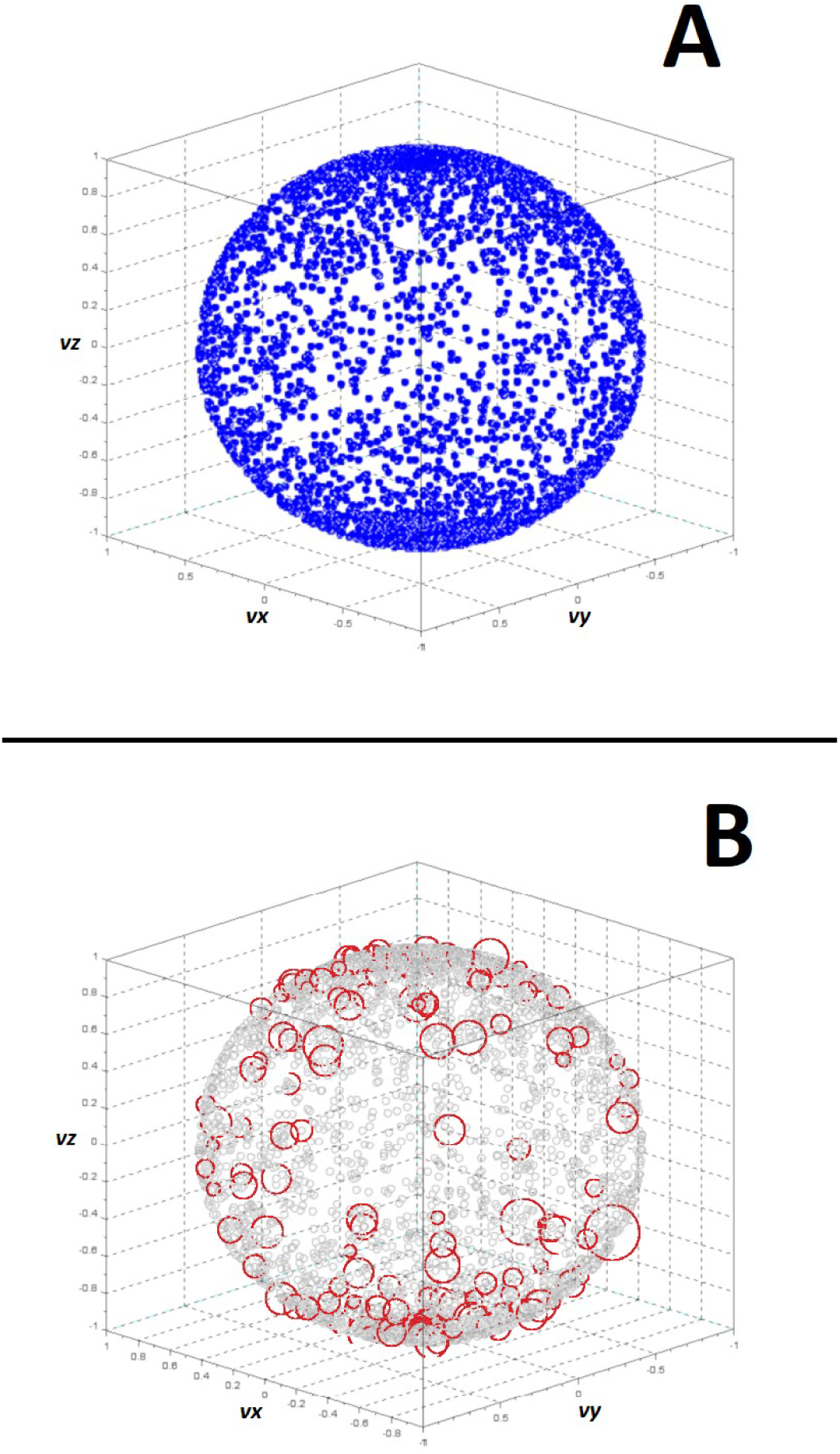
Ommatidia feature space projected to a spherical transformation. **A:** ommatidia cells (blue points) transformed to a spherical map and plotted by shape. **B:** cells for all row 8 ommatidia cells (red circles with a diameter relative to their pixel size) as identified in Supplemental File 2 plotted against the ommatidia cells (gray). One pixel equals 0.05 μm.

Figure 9A show the basic shape of and distribution of cells within this space, while Figure 9B demonstrates the location of all cells in ommatidia row 8 (see Supplemental File 2) embedded within this space. Figure 9B demonstrates that while anatomical spatial relationships are broken apart in spherical maps, angular relationships between cells are preserved. What is less clear is how to analyze these structures. Various types of topological data analysis (Wasserman, 2018) might be used in future studies to understand what non-trivial information can be extracted from these representations. Animations of subsampled spherical maps based on spatiotemporally-ordered strips of the differentiated eye imaginal disc might also be used to visually reveal information about the speed and acceleration of morphogenetic furrow progression.

## Discussion

A developmental process results in a number of spatial patterns. Some of these patterns describe differentiation as it unfolds. Other patterns reflect meta-features of the embryo at multiple spatial scales that might be biologically informative. One of our primary motivations comes from Proposition 250 in Gordon (1999). Proposition 250 states that the spacing patterns of cells (relative size and position) are indicative of contraction and expansion waves that occur during development. While more work needs to be done to make definitive statements about this proposition with respect to the *Drosophila* eye imaginal disc, we suspect that there is an interaction between the morphogenetic furrow and the relative location of differentiated cells (Courcoubetis et al., 2018).

We have proposed a series of quantitative approaches for understanding the interesting developmental properties of the *Drosophila* eye imaginal disc. We have also addressed our four computational hypotheses. The spherical map provides a uniform space where spatial variation due to artificial sources of curvature are removed. Further analysis of the data embedded in this structure is necessary to make more comprehensive statements about relationships between different regions of the ommatidia array.

While the ridge estimation procedure was used to yield labeled series for ommatidia rows, the analysis of the resulting graph might be the object of future work. As these ridge maps resemble fingerprint patterns, mathematical techniques for their analysis with broad application might be possible. Such an analysis also reveals new statistical features as well as local patterns in information content relevant to fluctuations in the developmental process. The independence of the ommatidia pattern from the wide range of cell sizes is particularly noteworthy.

There are a number of additional approaches that might be used in the future to uncover further developmental complexity. Our measurement of labeled ommatidia cell size resembles the ensemble averages of Torquato and Stillinger (2003) in their method to detect hyperuniformity of features in a spatial array. While we did not look for hyperuniformity amongst ommatidia structures, models of hyperuniformity and neural selection (Frankfort and Mardon, 2002) might be used to find additional patterns in these data. We can also look at other developmental systems (such as the development of horns in insects) for principles of developmental morphogenesis (Matsuda et al., 2017).

The *Drosophila* eye imaginal disc deserves further time lapse analysis, well beyond what we could do here with a single camera lucida drawing. For instance, the *shibire* mutant results in broad lines of no ommatidial formation when the temperature is raised, but ommatidia continue to form when it is lowered. If the temperature is raised, lowered, then raised and lowered again, two smooth broad lines are formed (Suzuki, 1974) (Figure 10). This suggests that the morphogenetic furrow, and perhaps differentiation waves in general, can be uncoupled from differentiation itself, and offers a powerful tool for investigating whether or not differentiation waves are the ultimate cause of cell differentiation, as has been proposed in Gordon (1999) and further explicated in Gordon and Gordon (2016).

**Figure 10.**
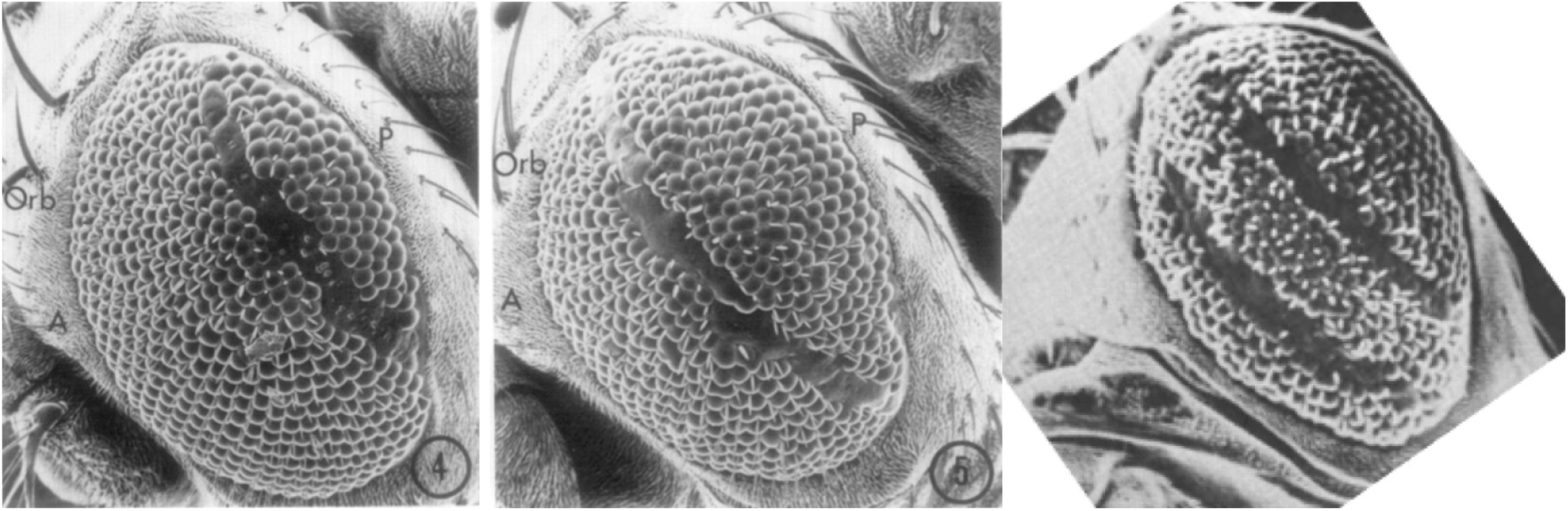
“The temperature sensitive mutant *shibire*^*ts*1^ of the fruit fly *Drosophila melanogaster* may be used to record the motion of the differentiation wave called the ‘morphogenetic furrow’ across the eye imaginal disc. **Left**: When the temperature is raised briefly from 22 to 29°C the wave keeps propagating, but the subsequent steps of differentiation of the cells into ommatidia fail. **Middle**: Here the temperature was raised briefly at a later time. The first two SEMs are from Poodry et al. (1973) with permission from Elsevier. **Right**: When the temperature is raised briefly twice, two lines of ommatidia are missing. The third SEM is rotated to the same orientation of the first two and is from (Suzuki, 1974) with permission of NRC Research Press” (Gordon and Gordon, 2016). Anterior is to the lower left, posterior to the upper right.

## Acknowledgements

We would like to highlight the contributions of Tanya Wolff and Kyle Harrington for their previous analytical efforts.

## Supplemental Files

**Supplemental File 1.** Digitized and then digitally retouched drawing of the *Drosophila* eye imaginal disc, hand corrected at 5840 pixels, adapted from the original drawing made by Tanya Wolff (Wolff, 1993), with image retouching done by Diana Gordon. The morphogenetic furrow propagates from posterior to anterior (left to right here) leaving the nascent ommatidia in its wake.

**Supplemental File 2.** Summary statistics (Counts, mean size, and mean position for all ommatidia labeled by their row and order from the posterior to the anterior side of the imaginal disc.

